# The language of mental images: Characterizing hippocampal contributions to imageable word use during event construction

**DOI:** 10.1101/2020.06.28.176131

**Authors:** Elizabeth Race, Camille Carlisle, Ruchi Tejwani, Mieke Verfaellie

## Abstract

Accumulating evidence suggests that the hippocampus plays a critical role in the creative and flexible use of language. For example, amnesic patients with hippocampal damage produce less coherent and cohesive verbal discourse when constructing narratives about the past, present, and future. A recent study by Hilverman and colleagues (2017) found that amnesic patients with hippocampal damage also use less imageable words during narrative construction compared to healthy controls. These results suggest that in addition to supporting language use at the discourse level, the hippocampus also influences the quality of language at the single word level. However, the generalizability of these results to different types of language production tasks and the relationship to patients’ broader impairments in episodic memory have yet to be examined. In the current study, we investigated whether amnesic patients with hippocampal damage produce less imageable words compared to healthy controls in two different types of language production tasks. In Experiment 1, participants constructed narratives about events depicted in visually presented pictures (picture narratives). In Experiment 2, participants constructed verbal narratives about remembered events from the past or simulated events in the future (past/future narratives). Across all types of narratives, patients produced words that were rated as having similar levels of imageability compared to controls. Importantly, this was the case both in patients’ picture narratives, which did not require generating details from long-term memory and were matched to controls’ with respect to narrative content, and in patients’ narratives about past/future events, which required generating details from long-term memory and which were reduced in narrative content compared to those of controls. These results reveal that the hippocampus is not necessary for the use of imageable representations at the linguistic level, and that hippocampal contributions to imageable word use are independent of hippocampal contributions to episodic memory.

## 1. Introduction

Accumulating evidence suggests that in addition to its pivotal role in episodic memory, the hippocampus also supports cognitive functions outside the domain of long-term memory, such as visual perception, attention, and short-term memory (Aly & Turk-Browne, 2018; Lee et al., 2005; Moscovitch, Cabeza, Winocur, & Nadel, 2016; Olsen, Moses, Riggs, & Ryan, 2012; Shohamy & Turk-Browne, 2013; Turk-Browne, 2019). Recently, there has been a growing interest in potential linguistic functions of the hippocampus (Corballis, 2019; Duff & Brown-Schmidt, 2012). Much of the evidence for hippocampal involvement in language has come from the study of language production in amnesic patients with hippocampal damage. Although linguistic functions are largely intact in these patients (Kensinger, Ullman, & Corkin, 2001; Milner, 1968; Skotko, Andrews, & Einstein, 2005), recent evidence suggests that hippocampal damage leads to qualitative changes in language production, particularly when constructing detailed and complex narratives that unfold over time (Duff, Hengst, Tranel, & Cohen, 2009; Hilverman, Cook, & Duff, 2016; MacKay, Burke, & Stewart, 1998; Race, Keane, & Verfaellie, 2015). For example, when describing personally experienced past events or imagined future events, amnesic patients with hippocampal damage produce narratives that are reduced in narrative cohesion and coherence (Caspari & Parkinson, 2000; Kurczek & Duff, 2011; MacKay et al., 1998; Race et al., 2015). Importantly, these discourse-level impairments are also present when patients construct detailed narratives based on visually presented pictures, in which the production of narrative content does not require retrieval from long-term memory and in which patients produce an equivalent amount of narrative detail as controls (Race et al., 2015; but see Keven, Kurczek, Rosenbaum, & Craver, 2018). Together with convergent neuroimaging data demonstrating hippocampal activity during language tasks (Piai et al., 2016), such results suggest that functions supported by the hippocampus, such as relational binding and on-line integration, support the constructive and flexible use of both mnemonic and linguistic representations (Duff & Brown-Schmidt, 2012; MacKay, James, Hadley, & Fogler, 2011; Olsen et al., 2012).

The results of a recent neuropsychological study by Hilverman and colleagues (2017) suggest that the hippocampus may also play a role in language production at the level of single words (see also Park, St-Laurent, McAndrews, & Moscovitch, 2011). In this study, amnesic patients with damage to the hippocampus were cued to produce narratives about specific moments in time during the lived past, imagined past, imagined present, and imagined future. Consistent with the broader literature, patients’ narratives contained fewer words and episodic details compared to those of controls (e.g., Hassabis, Kumaran, Vann, & Maguire, 2007; Kurczek et al., 2015; Race, Keane, & Verfaellie, 2011). The authors also measured the quality of individual words (i.e., word imageability) and found that, compared to controls, patients used words that were rated lower in imageability. Word imageability is a measure of the degree to which a word evokes a mental image or generates an internal visual representation (Paivio, Yuille, & Madigan, 1968). For example, the word “eagle” is rated higher in imageability than the word “trust.” Imageability is thought to reflect the activation of visual-perceptual knowledge and is typically associated with neural activity in anterior and inferior temporal lobe regions involved in the storage of semantic information and object recognition (e.g., Bonner et al., 2009; Loiselle et al., 2012; Yi, Moore, & Grossman, 2007). However, recent neuroimaging studies have found that stimulus imageability also modulates activity in the hippocampus and medial temporal lobe (MTL) cortex, both in memory tasks (e.g., Caplan & Madan, 2016; Klaver et al., 2005) and in non-memory tasks such as word association and semantic similarity (Bonner, Price, Peelle, & Grossman, 2016; Sabsevitz, Medler, Seidenberg, & Binder, 2005; Wise et al., 2000). The lesion findings by Hilverman and colleagues (2017) align with these results and suggest that the hippocampus may play a critical role in the verbal description of imageable representations.

Although the results by Hilverman and colleagues (2017) suggest that the hippocampus may play a broader role in language production at the single word level, several important outstanding questions remain. First, it is currently unclear whether hippocampal contributions to imageable word use represent a novel linguistic function of the hippocampus or simply reflect the canonical role of the hippocampus in episodic memory. It is well established that the (re)constructive processes that support the generation of past and future scenarios, such as the retrieval and/or recombination of details stored in long-term memory, critically depend on the hippocampus (Schacter, 2012; Schacter and Addis, 2007). It is important to note that in the study by Hilverman and colleagues (2017), imageability impairments were observed in amnesia in the context of narrative tasks that required the retrieval and (re)combination of narrative content from memory and in which amnesic patients demonstrated concurrent reductions in narrative content and detail (Kurczek et al., 2015). That is, in addition to producing words that were less imageable, patients also produced significantly *fewer* words and episodic details in their narratives. Although Hilverman and colleagues covaried for word count and proportion of internal details in their analyses, the contribution of the hippocampus to word use in these narratives could still be mediated by its established role in episodic memory. Second, it is currently unclear whether the reduction in imageable word use in amnesia is a generalizable characteristic of amnesic patients’ language production, or whether this deficit is specific to certain types of verbal narratives. In the study by Hilverman and colleagues (2017), participants were instructed to select a specific moment from an event and to “produce a narrative of the setting and experience of that specific moment,” which likely encouraged use of static, scene-based imagery. In contrast, many of our everyday verbal narratives involve telling stories and describing events that unfold across space and time to communicate our feelings, plans, and ideas (Corballis, 2013; Suzuki, Feliu-Mojer, Hasson, Yehuda, & Zarate, 2018). An outstanding question is whether the role of the hippocampus in imageable word use generalizes to the construction of these more naturalistic, event-based narratives.

To investigate these questions, the current study analyzed the imageability of words produced in four different types of verbal narratives that were previously acquired from amnesic patients with hippocampal damage and healthy controls. In Experiment 1, we investigated the imageability of words produced during the description of visually presented pictures (picture description). Picture description provides a constrained method of eliciting verbal narratives and has been widely used to assess the language abilities of clinical populations (Bird, Lambon Ralph, Patterson, & Hodges, 2000). Participants produced two different picture description narratives: (1) static and (2) event-based. For static narratives, participants were shown a picture of a scene and were instructed to describe what they saw in the picture. For event-based narratives, participants were shown a picture of a scene and were instructed to tell a *story* about the events depicted in the picture. Importantly, both types of picture narratives place low demands on retrieval from long-term memory (i.e., narrative content is provided by the picture and does not have to be generated from memory) and we have previously demonstrated that the amount of narrative content produced by amnesic patients and controls in these narratives is matched (Race et al., 2011; Race, Keane, & Verfaellie, 2013). This enabled us to dissociate potential impairments in imageable word use from patients’ impairments in long-term memory. If the contribution of the hippocampus to the production of imageable words reflects a general linguistic function outside of its role in memory, imageability performance in amnesia should be impaired in these picture tasks. Alternatively, if the contribution of the hippocampus to imageable word use is mediated by its role in long-term memory, imageability performance in amnesia should be intact in these narratives. A final possibility is that patients might produce less imageable words in only one type of narrative task. In prior neuroimaging studies, imageability effects in the hippocampus during non-memory tasks have been shown to depend on the nature of the task demands (e.g., present for semantic similarity judgements but not for lexical decisions; Binder, Westbury, McKiernan, Possing, & Medler, 2005; Sabsevitz et al., 2005). Given that Hilverman and colleagues (2017) observed that patients used less imageable words in narratives that encouraged static imagery, deficits in imageable word use may be particularly pronounced in patients’ static picture descriptions. To foreshadow our results, we found that imageable word use was intact in amnesic patients in both static and event-based picture narratives. These results argue against a broader role of the hippocampus in language production at the single word level when controlling for demands on memory retrieval, and instead suggest that hippocampal contributions to imageable word use may be mediated by the role of the hippocampus in episodic memory.

Experiment 2 further investigated the relationship between imageable word use and episodic memory by testing the scope of imageability deficits in amnesia when constructing event narratives about the lived past and the imagined future. Like the narratives in the study by Hilverman and colleagues (2017), these narratives require retrieving event details from long-term memory and have been shown to be less detailed in amnesic patients with damage to the hippocampus (Race et al., 2011). However, whereas the narratives in the study by Hilverman and colleagues (2017) involved describing details from a snapshot of an event that is constrained in time and space, these future/past narratives involved describing a series of events that unfolded over space and time. This allowed us to investigate whether previously observed deficits in imageable word use in amnesia generalize to verbal narratives about more dynamic, unfolding events that draw upon episodic memory. If the hippocampus contributes to the production of imageable words through its role in episodic memory, deficits in imageable word use in amnesia should generalize to these future/past narratives.

## Experiment 1: Picture Narratives

### 2. Materials and Methods

#### 2.1 Participants

Nine amnesic patients with MTL lesions participated in the study. All of the amnesic patients participated in a prior study investigating discourse cohesion and coherence in narratives about novel future events, experienced past events, and events in pictures (Race et al., 2015). In addition, eight of the amnesic patients participated in a prior study investigating the nature of the content of these narratives (Race et al., 2011). Neuropsychological profiles for the patients are described in Table 1 and indicate severe impairments isolated to the domain of memory with profound deficits in new learning. Volumetric data for the hippocampus and MTL cortices were available for five patients. Two of the anoxic patients (P05 and P09) had damage limited to the hippocampus, and two of the encephalitic patients (P01 and P02) and one of the anoxic patients (P04) had damage to the hippocampus and surrounding parahippocampal gyrus. No common volume reductions were found outside the hippocampus. MRI could not be obtained for the remaining patients because of medical contraindications. For the encephalitic patient P06, a computerized tomography (CT) scan was available and visual inspection indicated extensive hippocampal and parahippocampal gyrus damage. For the remaining patients, MTL pathology can be inferred on the basis of etiology and neuropsychological profile.

**Table 1.** Patient Demographic, Neuropsychological and Neurological Characteristics

| Patient | Etiology | Age | Edu | WAIS | WMS, III |  |  | Hipp | Subhipp |
| --- | --- | --- | --- | --- | --- | --- | --- | --- | --- |
|  |  |  |  | III<br>VIQ | GM | VD | AD | Vol Loss | Vol Loss |
| P01 | Encephalitis | 55 | 14 | 92 | 45 | 56 | 55 | 73% | 78%* |
| P02 | Encephalitis | 66 | 12 | 106 | 69 | 68 | 77 | 66% | 72%+ |
| P03 | Anoxia/ischemia | 60 | 12 | 83 | 52 | 56 | 55 | N/A | N/A |
| P04 | Anoxia + left<br>temporal lobectomy | 46 | 16 | 86 | 49 | 53 | 52 | 63% | 60%^ |
| P05 | CO poisoning | 54 | 14 | 111 | 59 | 72 | 52 | 22% | - |
| P06 | Encephalitis | 82 | 18 | 135 | 45 | 53 | 58 | N/A | N/A |
| P07 | Cardiac arrest | 58 | 17 | 134 | 70 | 75 | 67 | N/A | N/A |
| P08 | Cardiac arrest | 60 | 16 | 110 | 62 | 68 | 61 | N/A | N/A |
| P09 | Stroke | 55 | 18 | 119 | 67 | 75 | 55 | 58% | - |

*Note.* Age = Age (years); Edu = Education (years); WAIS, III = Wechsler Adult Intelligence Scale, III; VIQ = Verbal IQ; WMS, III = Wechsler Memory Scale, III; GM = General Memory; VD = Visual Delayed; AD = Auditory Delayed; WM = Working Memory; Hipp Vol Loss = Bilateral Hippocampal Volume Loss; Subhipp Vol Loss = Parahippocampal Gyrus Volume Loss; CO = carbon monoxide
\* = volume loss in bilateral anterior parahippocampal gyrus and left posterior parahippocampal gyrus.
+ = volume loss in bilateral anterior parahippocampal gyrus and right posterior parahippocampal gyrus.
^ = volume loss in left anterior parahippocampal gyrus.

Twelve healthy controls also participated, all of whom had participated in the prior studies by Race et al. (2011; 2015). The control subjects were matched to the patient group in terms of mean age (60 ± 12.2 years), education (14 ± 2.0 years), and verbal IQ (105 ± 15.7). As reported by Race et al. (2011, 2013), quantitative assessment revealed that the patients’ descriptions of the future and past contained fewer episodic details than those of controls, whereas their picture descriptions contained an equivalent number of episodic details compared to those of controls. This pattern of impairment was also present in the additional amnesic patient included in the present study (P09), who provided fewer episodic details than controls in his future and past narratives (z scores < −2) but did not provide fewer episodic details in his picture narratives (z scores > .9). All participants were paid for their participation and provided informed consent in accordance with the procedures of the Institutional Review Boards at Boston University and the VA Boston Healthcare System.

#### 2.2 Stimuli

This study is a reanalysis of the picture description data reported by Race et al. (2011; 2013). Participants were shown detailed drawings of scenes, one at a time, that depicted characters engaged in various activities (e.g., a picnic at a park). In the “static” condition, participants were instructed to describe what they saw in the picture in as much detail as possible without creating a story about the picture (Race et al., 2013). In the “dynamic” condition, participants were instructed to imagine that the picture was a scene taken from a movie and to tell a story about what was going on in the scene (Race et al., 2011). Five narratives in each condition were audiotaped and transcribed for analysis.

#### 2.3 Scoring

Narratives were scored following the procedures used by Hilverman et al. (2017). Words in each narrative were first coded as either a function word or a content word. Content words were defined as adjectives, adverbs, nouns, and verbs. Function words included pronouns (i.e. she, me, it, what, etc.), auxiliary verbs (i.e. do, have, been, etc.), articles (i.e. the, an, etc.), particles (i.e. well, um, etc.), and conjunctions (i.e. and, but, if, etc.) The MRC Psycholinguistic database was used to score each unique content word for verbal frequency and imageability. More information on these measures can be found on the MRC Psycholinguistic database website (http://websites.psychology.uwa.edu/school/MRCDatabase/uwa_mrc.htm). Contractions were manually changed to non-contracted words before scoring. Words not included in the MRC database were not scored. The percentage of unique content words scored in each type of narrative did not differ for patients (*M* = 50%) and controls (*M* = 51%) (*t*(19) = 2.19, *p* = .31).

### 3. Results

#### 3.1 Word Count

Healthy control participants produced an average of 205 total words (*SD* = 95) per narrative and amnesic patients produced an average of 158 total words (*SD* = 90) per narrative. Total word count was entered into a two-way mixed factorial ANOVA with factors of group (patient, control) and picture narrative condition (static, dynamic). There was no main effect of group (*F*(1,19) = 1.46, *p* = .24, *η*_p2_ = .07), indicating that the number of words produced by patients and controls in the picture narrative tasks did not significantly differ. There was also no main effect of picture narrative condition (*F(* 1,19) = 3.10, *p* = .09, *η*_p2_ = .14) nor group x condition interaction (*F*(1,19) = .43, *p* = .52, *η*_p2_ = .02). We next analyzed whether there was any difference in the number of unique content words used by each group of participants. Healthy control participants produced an average of 69 unique content words (*SD* = 24) and amnesic patients produced an average of 56 unique content words (*SD* = 28) per narrative. A two-way mixed factorial ANOVA with factors of group (patient, control) and narrative condition (static, dynamic) revealed that there was no main effect of group (*F*(1,19) = 1.37, *p* = .26, *η*_p2_ = .07). There was also no main effect of picture narrative condition (*F*(1,19) = 1.68, *p* = .21, *η*_p2_ = .08) nor group x condition interaction (*F*(1,19) = .03, *p* = .87, *η*_p2_ = .002).

#### 3.2 Verbal Frequency

The unique content words produced by healthy controls and amnesic patients were similar in terms of mean verbal frequency (controls: *M* = 217, *SD* = 42; patients: *M* = 231, *SD* = 51). When verbal frequency ratings were entered into a two-way mixed factorial ANOVA with factors of group (patient, control) and picture narrative condition (static, dynamic), there was no main effect of group (*F*(1,19) = .96, *p* = .34, *η*_p2_ = .05). There was also no main effect of condition (*F*(1,19) = 3.66, *p* = .07, *η*_p2_ = .16) nor group x condition interaction (*F*(1,19) = .008, *p* = .93, *η*_p2_ < .001).

#### 3.3 Imageability

Our primary analysis of interest concerned the imageability ratings of the unique words produced by amnesic patients and healthy controls (**Figure 1A**). The average imageability ratings were remarkably similar across groups (controls: *M* = 450, *SD* = 35; patients: *M* = 452, *SD* = 23). Word imageability ratings were entered into a two-way mixed factorial ANOVA with factors of group (patient, control) and narrative condition (static, dynamic). A main effect of condition (*F*(1,19) = 19.79, *p* < .001, *η*_p2_ = .51) reflected that words produced in participants’ static narratives were more imageable than words produced in participants’ dynamic narratives. However, there was no main effect of group (*F*(1,19) = .03, *p* = .86, *η*_p2_ = .002) nor group x time period interaction (*F*(1,19) = 1.85, *p* = .19, *η*_p2_ = .09), indicating that the imageability of the words used in patients’ and controls’ picture narratives did not significantly differ. Data from the static and dynamic narratives were also analyzed using a one-way ANCOVA with word count and verbal frequency as covariates. When adjusting for word count and verbal frequency, there was still no significant main effect of group for either static picture narratives (*F*(1,17) = .31, *p* = .58, *η*_p2_ = .02) nor dynamic picture narratives (*F*(1,17) = 3.96, *p* = .06, *η*_p2_ = .19). In fact, post-hoc analysis of the adjusted marginal means revealed that patients used words that had numerically higher imageability ratings (*M* = 444) in their dynamic narratives compared to controls (*M* = 430). The finding that patients and controls use similarly imageable words in their picture narratives was confirmed on an individual basis when looking at patients’ z-scores: None of the individual patients demonstrated significantly impaired performance compared to controls as defined by a z-score cut-off of −1.96 (z score range: −1.49 to 1.52).

**Figure 1.**
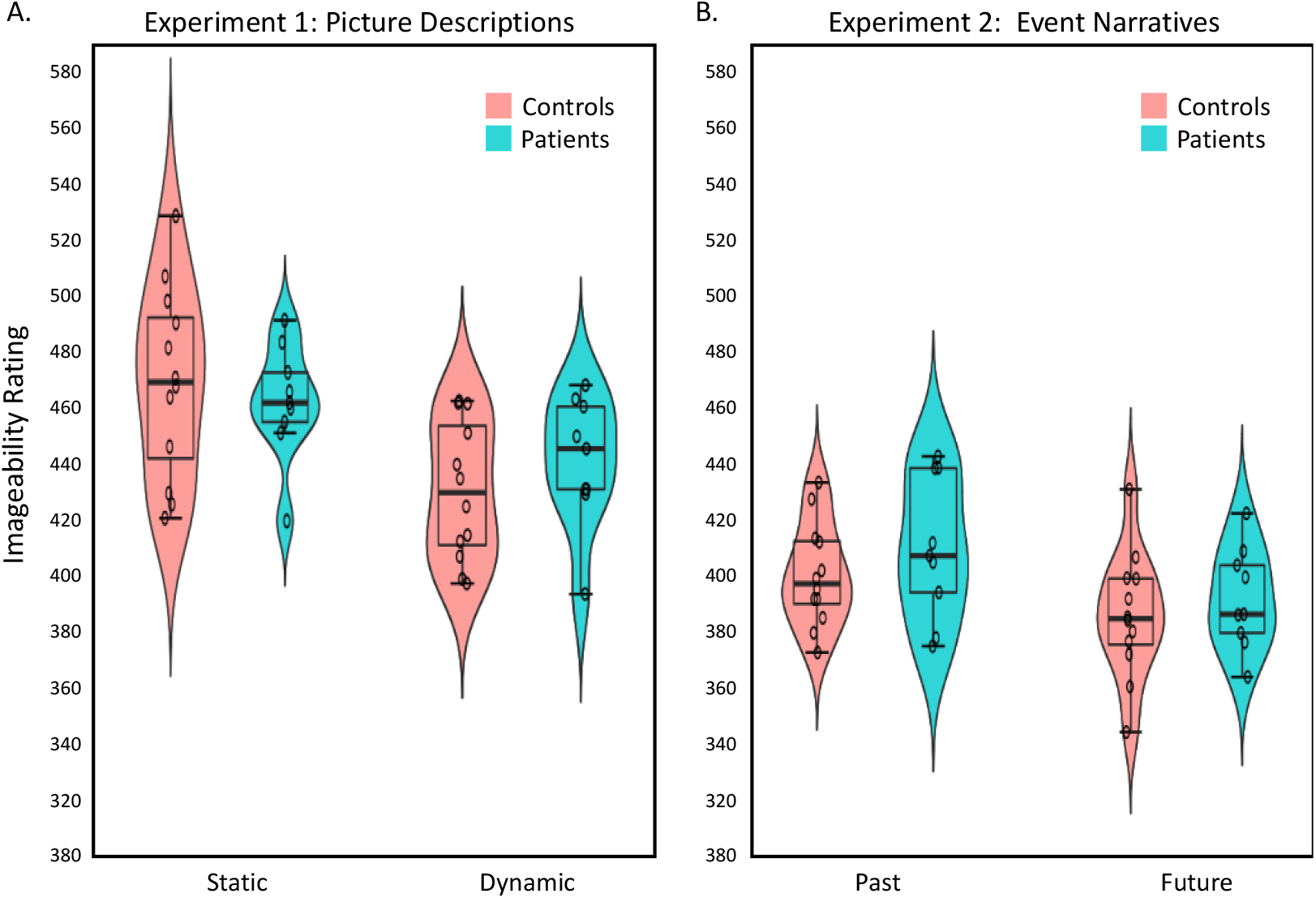
Imageability Ratings in Experiments 1 and 2. Average imageability ratings of words produced in narratives about (A) past and future events and (B) static and event-based picture descriptions. Healthy controls (pink) and amnesic patients (blue) produced individual words rated similarly in imageability in all types of narratives. Boxplots depict median value, interquartile range, and minimum/maximum values. Violin plot depicts smoothed distribution curve. Circles represent individual participant data within each group.

#### 3.4 Relationship to Episodic Memory

We next explored whether the imageability ratings of the words produced in patients’ picture narratives were related to the extent of patients’ deficit in episodic memory as measured by neuropsychological assessment. For dynamic narratives, there was no correlation between patients’ imageability ratings and their scores on the Visual Delay (*r* = .03, *p* = .94) or General Memory (*r* = −.33, *p* = .39) subscales of the Wechsler Memory Scale, III. For static narratives, although the correlations did not reach statistical significance, there appeared to be a positive relationship between patients’ imageability ratings and their scores on (a) the Visual Delay subscale of the Wechsler Memory Scale, III (*r* = .58, *p* = .10) and (b) the General Memory subscale of the Wechsler Memory Scale, III (*r* = .50, *p* = .17).

#### 3.5 Distribution of Imageability Scores

To examine whether the distribution of imageable words used by patients and controls differed, we performed an additional analysis of the number of words used by each group in each quintile of imageability ratings (**Supplementary Table 1**). The range of imageability ratings was first calculated across healthy controls, separately for static narratives and for dynamic narratives, in order to define five quintile bins for each type of picture narrative. The number of unique words in each quintile bin was then counted in each participant and the distributions were compared in patients and controls using a Chi-Square test. For static narratives, the distribution of words across imageability bins did not differ between patients and controls (*X*_2_ (2, *N* = 21) = .12, *p* = .99). For dynamic narratives, the distribution of words across imageability bins also did not differ between patients and controls (*X*_2_ (2, *N* = 21) = .81,*p* = .94).

### 4. Interim Discussion

The results from Experiment 1 demonstrate that when constructing narratives about visually presented pictures, patients with amnesia use words that are similar in imageability to those used by controls. Importantly, these narratives did not require retrieval of narrative content from long-term memory and were matched in terms of narrative content across patients and controls. These findings suggest that previously observed reductions in imageable word use in amnesia (i.e., Hilverman et al., 2017) may reflect hippocampal contributions to episodic memory rather than language production per se. Experiment 2 aimed to further test this hypothesis by investigating patients’ word use when constructing narratives that draw upon episodic memory and have been shown to be impoverished in amnesia (past/future narratives). In contrast to the narratives in the study by Hilverman and colleagues (2017), in which participants described a snapshot of time within an event, we examined dynamic, event-based narratives in which participants described events that unfolded over time. This allowed us to investigate whether previously observed deficits in imageable word use in amnesia generalize to dynamic, event-based verbal narratives about the past and future.

## Experiment 2: Future and Past Narratives

### 5. Materials and Methods

#### 5.1 Participants

The same participants were tested in Experiments 1 and 2.

#### 5.2 Stimuli

This study is a reanalysis of past/future narrative data reported by Race et al. (2011; 2015). Narratives about the past and the future were generated by having participants either recall specific personal events about the past (e.g., graduation ceremony) or imagine specific personal events about the future (e.g., winning the lottery). Participants were given three minutes to describe the event in as much detail as possible, and were instructed to describe where and when the event is taking place, who is there, how they feel, and what they are thinking. Within the allotted three minutes, participants continued with their descriptions without interference from the examiner until they came to a natural ending point. Narratives were audiotaped and transcribed for analysis. In the current study, two narratives about the future and two narratives about the past were selected from the larger sample for analysis (mirroring the procedures used by Race et al., 2015).

#### 5.3 Scoring

Narratives were scored following the same procedures used in Experiment 1. The percentage of unique content words scored in each type of narrative did not differ for patients and controls in narratives about the past (*M* = 68% and 67%, respectively) or narratives about the future (*M* = 64% and 66%, respectively) (*t*s(19) < .44, *p*s > .66).

### 6. Results

#### 6.1 Word Count

Healthy control participants produced an average of 125 words (*SD* = 50) per narrative and amnesic patients produced an average of 78 words (*SD* = 47) per narrative. Total word count was entered into a two-way mixed factorial ANOVA with factors of group (patient, control) and time period (past, future). There was a main effect of group (*F*(1,19) = 6.09, *p* = .02, *η*_p2_ = .24), indicating that patients produced overall fewer words than controls, but no main effect of time period (*F*(1,19) = .009, *p* = .93, *η*_p2_ = .001) nor group x time period interaction (*F*(1,19) = .05, *p* = .83, *η*_p2_ = .002). We next analyzed whether there was any difference in the number of unique content words used by each participant group, again using a two-way mixed factorial ANOVA with factors of group (patient, control) and time period (past, future). There was a main effect of group (*F*(1,19) = 6.62, *p* = .02, *η*_p2_ = .26), reflecting that patients produced fewer unique content words than controls, but no main effect of time period (*F*(1,19) = .17, *p* = .68, *η*_p2_ = .009) nor group x time period interaction (*F*(1,19) = .02, *p* = .89, *η*_p2_ = .001). Together, these results converge with the prior finding that amnesic patients produce narratives about the past and future that contain fewer episodic details compared to those of controls (Race et al., 2011).

#### 6.2 Verbal Frequency

The mean verbal frequency of the unique content words produced by healthy controls and amnesic patients in their future/past narratives were similar (controls: *M* = 291, *SD* = 41; patients: *M* = 275, *SD* = 78). When verbal frequency ratings were entered into a two-way mixed factorial ANOVA with factors of group (patient, control) and picture narrative condition (future, past), there was no main effect of group (*F*(1,19) = .82, *p* = .38, *η*_p2_ = .04). There was also no main effect of condition (*F*(1,19) = .47, *p* = .50, *η*_p2_ = .02) nor group x condition interaction (*F*(1,19) = 2.08, *p* = .17, *η*_p2_ = .10).

#### 6.3 Imageability

The average imageability rating of content words produced in narratives about the past and future was very similar in healthy controls (*M* = 392; *SD* = 22) and amnesic patients (*M* = 400; *SD* = 24) (**Figure 1B**). When data were entered into a two-way mixed factorial ANOVA with factors of group (patient, control) and time period (past, future), there was a main effect of time period (*F*(1,19) = 9.33, *p* = .007, *η*_p2_ = .33), indicating that words produced in past narratives were more imageable than words produced in future narratives, similar to the findings reported by Hilverman and colleagues (2017). However, there was no main effect of group (*F*(1,19) = 1.10, *p* = .31, *η*_p2_ = .06) nor group x time period interaction (*F*(1,19) = .12, *p* = .73, *η*_p2_ = .006), indicating that the words produced in patients’ and controls’ narratives were similarly imageable. Data from the past and future narratives were also analyzed using a one-way ANCOVA with word count, verbal frequency, and proportion of episodic details as covariates. There was no main effect of group when adjusting for word count, verbal frequency, and proportion of episodic features for narratives about past events (*F*(1,16) = 1.47, *p* = .24, *η*_p2_ = .08) or narratives about future events (*F*(1,16) = 3.53, *p* = .08, *η*_p2_ = .18). For future events, the trend towards a difference in adjusted imageability ratings between patient and controls reflected numerically *greater* ratings in patients (*M* = 397.52) compared to controls (*M* = 379.28). Covariate-adjusted imageability ratings (studentized residuals) were also greater for past compared to future narratives (*F*(1,19) = 9.33,*p* < .01, *η*_p2_ = .33). The finding that patients and controls use similarly imageable words in their narratives about the past and future was confirmed on an individual basis when looking at patients’ z-scores: None of the individual patients demonstrated significantly impaired performance compared to controls as defined by a z-score cut-off of −1.96 (z score range: −1.45 to +2.30).

#### 6.4 Relationship to Episodic Memory

We next explored whether the imageability ratings of the words produced in patients’ narratives about the past or future were related to the extent of patients’ deficit in episodic memory as measured by neuropsychological assessment. No correlation was found between imageability ratings in patients’ past narratives and their scores on the Visual Delay subscale (*r* = −.03, *p* = .94) or General Memory subscale (*r* = −.05, *p* = .88) of the Wechsler Memory Scale, III. Similarly, no correlation was found between imageability ratings in patients’ future narratives and their scores on the Visual Delay subscale (*r* = −.25, *p* = .51) or General Memory subscale (*r* = −.17, *p* = .67) of the Wechsler Memory Scale. Furthermore, there was also no correlation between imageability ratings in patients’ past or future narratives and the proportion of episodic details produced in these narratives (past: *r* = .17, *p* = .47; future: *r* = .42, *p* = .06).

#### 6.5 Distribution of Imageability Scores

To examine whether the distribution of imageable words used by patients and controls differed, we performed an additional analysis of the number of words used by each group in each quintile of imageability ratings (**Supplementary Table 1**). The range of imageability ratings was first calculated across healthy control participants, separately for past and for future narratives, in order to define five quintile bins for each type of narrative. The number of unique words in each quintile bin was then counted in each participant and the distributions were compared in patients and controls using a Chi-Square test. For past narratives, the distribution of words across imageability bins did not differ between patients and controls (*X*_2_ (2, *N* = 21) = .19, *p* = .99). For future narratives, the distribution of words across imageability bins also did not differ between patients and controls (*X*_2_ (2, *N* = 21) = .30, *p* = .99).

### 7. General Discussion

The current study investigated whether the hippocampus plays a critical role in imageable word use during narrative construction. In four different types of verbal narratives, amnesic patients with hippocampal lesions were able to generate imageable words as well as controls. This was the case both for (1) picture narratives, which do not require mentally generating details from long-term memory and are matched across groups in terms of narrative content, and (2) narratives about the past and future, which depend on access to long-term memory representations, and for which patients generate fewer details than controls. These results distinguish between the quantity and quality of individual linguistic details produced in amnesia during narrative construction, and suggest that the use of imageable linguistic representations can be supported by regions outside the hippocampus.

Although not included in traditional models of language, accumulating evidence suggests that the hippocampus contributes to the online use and processing of language, especially at the discourse level when linguistic elements must be flexibly integrated and maintained (Duff & Brown-Schmidt, 2012; Duff, Gupta, Hengst, Tranel, & Cohen, 2011; Kurczek, Brown-Schmidt, & Duff, 2013; Piai et al., 2016; Race et al., 2015). The contributions of the hippocampus to language use and processing have been attributed to its role in relational binding and the online integration of mnemonic and linguistic representations. However, a recent observation that amnesic patients with hippocampal damage produce less imageable words during narrative construction suggests that the hippocampus also supports qualitative aspects of language use at the single word level (Hilverman et al., 2017). The current study addressed an important outstanding question: whether hippocampal contributions to imageable word use represent a novel linguistic function of the hippocampus or instead reflect the canonical role of the hippocampus in episodic memory. Prior work could not distinguish between these possibilities given that deficits in imageable word use in amnesia always occurred in the context of concurrent deficits in narrative content. The observation in Experiment 1 that amnesic patients and controls produce similarly imageable words in two different picture description tasks that do not require retrieval from long-term memory reveals that the hippocampus is not critical for the access or use of imageable representations at the linguistic level. This aligns with prior work indicating that medial temporal lobe damage does not affect the use of lexical representations or grammatical processing (e.g., Kensinger et al., 2001) and reveals that the hippocampus is not necessary for imageable word use in non-mnemonic tasks.

Experiment 2 was conducted to examine whether the hippocampus plays a critical role in imageable word use in the context of narrative tasks that require retrieving details from long-term memory. Hilverman and colleagues (2017) previously proposed that the hippocampus contributes to imageable word use through its role in declarative memory and suggested that the use of less imageable words in amnesia directly relates to patients’ impoverished long-term memory representations. If this is the case, then imageable word use should be impaired in amnesia in the context of tasks that draw upon episodic memory. The findings of Experiment 2 do not support this proposal: Despite producing fewer details in their narratives about the past and future, amnesic patients produced individual narrative details that were as imageable as those generated by controls. In addition, no relationship was observed between the imageability of the words used in patients’ narratives and the extent of their episodic memory deficits as measured by neuropsychological assessment. The dissociation between the quantity and quality of the individual details produced in patients’ narratives suggests that the level of detail comprising one’s simulations of the past and future does not always affect the language used to describe these simulations. This observation is particularly interesting given that patients’ verbal narratives about constructed events are typically rated as being qualitatively inferior (e.g., less vivid) compared to those constructed by controls (e.g., Hassabis et al., 2007; Kurczek et al., 2015). This raises the intriguing possibility that qualitative ratings of event narratives may be more closely related to overall features of the narrative, such as the amount of narrative detail or discourse-level features of continuity and contextual organization, rather than the quality of the individual words used to convey the narrative. Indeed, in prior work we demonstrated that amnesic patients produce narratives that are rated lower in measures of narrative coherence and cohesion (Race et al., 2015), both in the context of narratives that draw upon episodic memory (future/past narratives) and in the context of narratives that do not require retrieval from episodic memory (picture narratives). More broadly, the results of Experiment 2 inform the prior results of Hilverman and colleagues (2017) and suggest that the hippocampus does not contribute to imageable word use through its role in episodic memory.

Why, then, might hippocampal lesions impair imageable word use in some contexts but not in others? Methodological differences between the present study and the study by Hilverman and colleagues (2017) may provide some insight into this question. In the Hilverman study, participants provided a one-to two-minute overview of an event and then constructed a narrative about the setting and experience of a specific moment within that event. Importantly, imageability ratings were only collected for words produced during the latter narrative when participants described a snapshot of time within the larger event. In contrast, participants in the present study were instructed to construct event-based future and past narratives that unfolded more dynamically over time. This difference in the spatiotemporal context of the narrative (e.g., static vs. dynamic) likely influenced the degree to which participants used spatial imagery or spatial context during event construction and could have influenced the nature or phenomenology of retrieved representations or the constructed events more broadly. For example, it is known that the spatial features of retrieval cues can influence the vividness of remembered or imagined scenarios (Robin & Moscovitch, 2014, Hebscher, Levine, & Gilboa, 2018; Robin, Wynn, & Moscovitch, 2016; Sheldon & Chu, 2017) and that phenomenological aspects of memory retrieval are also influenced by the degree to which the layout of a scene is instantiated in memory (Rubin, Deffler, & Umanath, 2019). A recent study by Sheldon and colleagues (2019) directly compared the effects of emphasizing spatial versus action-based contexts prior to generating past and future autobiographical events, and found that emphasizing spatial context led to the generation of a greater proportion of perception-based details (e.g., sensory features of objects or spatial contextual elements) compared to emphasizing action-based details. Further, spatiotemporally specific cues might place higher demands on the integration of perceptual details, which has been proposed to influence the recruitment of the hippocampus during mental construction (Sheldon & Levine, 2016). Describing the details of a specific spatiotemporal event might also encourage more precise scene-based imagery, which has been associated with hippocampal function during memory and mental simulation (Hassabis & Maguire, 2007, St-Laurent, Moscovitch, Jadd, & McAndrews, 2014; Yonelinas, 2013, Cowell, Barense, & Sadil, 2019; Sheldon & Levine, 2016, Bird, Bisby, & Burgess, 2012, but see Kim et al., 2013). This proposal aligns with theories that emphasize spatiotemporal coding as a primary feature of hippocampal representation and coding (e.g., Byrne, Becker, & Burgess, 2007; Ekstrom & Ranganath, 2018; Ekstrom & Yonelinas, 2020; Hasselmo, Hinman, Dannenberg, & Stern, 2017; Turk-Browne, 2019; Robin, Buchsbaum, and Moscovitch, 2018). Indeed, increased activity in the hippocampus has been observed when remembering or imagining more spatiotemporally specific events (Addis, Cheng, Roberts, & Schacter, 2011; Palombo, Hayes, Peterson, Keane, & Verfaellie, 2018) and hippocampal activation has been shown to increase parametrically with the spatial specificity of an imagined scene (Bird, Capponi, King, Doeller, & Burgess, 2010). Future research should more directly test whether hippocampal contributions to the retrieval of imageable mnemonic representations during discourse depends on the spatiotemporal specificity of the narrative cues.

The presence of imageability impairments in amnesia may also depend on the extent and location of patients’ lesions. Recent theoretical models of hippocampal function have emphasized the functional heterogeneity of the hippocampus (Dalton, Zeidman, McCormick, & Maguire, 2018; Nadel, Hoscheidt, & Ryan, 2013; Poppenk, Evensmoen, Moscovitch, & Nadel, 2013; Strange, Witter, Lein, & Moser, 2014; Zeidman & Maguire, 2016). For example, functional subdivisions have been proposed along the anterior-posterior axis of the hippocampus, with differential involvement along the anterior-posterior axis in more conceptual versus perceptual representation (Sheldon, Fenerci, & Gurguryan, 2019; Sheldon & Levine, 2016), coarser versus fine-grained representation (Poppenk et al., 2013), and scene-based versus more general constructive imagery (Dalton et al., 2018). Hippocampal lesions that differ with respect to their anterior-posterior extent might therefore differentially affect the representation or access to more imageable information in memory. Although the precise location of patients’ lesions within subregions of the hippocampus could not be differentiated in the current study, future studies using higher-resolution structural or functional mapping could investigate whether specific regions along the anterior-posterior axis of the hippocampus are particularly important for the retrieval or representation of mental images during event construction.

In conclusion, the present neuropsychological results reveal that the hippocampus does not play an obligatory role in the use of imageable words during narrative construction, and that imageable word use can be intact following hippocampal lesions even in the face of severe deficits in episodic memory. More broadly, the present study adds to growing body of work investigating non-mnemonic functions of the hippocampus and reveals that whereas the hippocampus plays a critical role in some forms of linguistic processing, it is not always critical for qualitative aspects of language use at the single word level. The present results also suggest promising avenues for future work to further specify under what circumstances hippocampal processes or representations support language use and processing.

## Acknowledgements

The authors thank Dr. Margaret Keane for helpful discussions about research design, data analysis, and interpretation of results.

## Supplement

**Supplementary Table 1.**
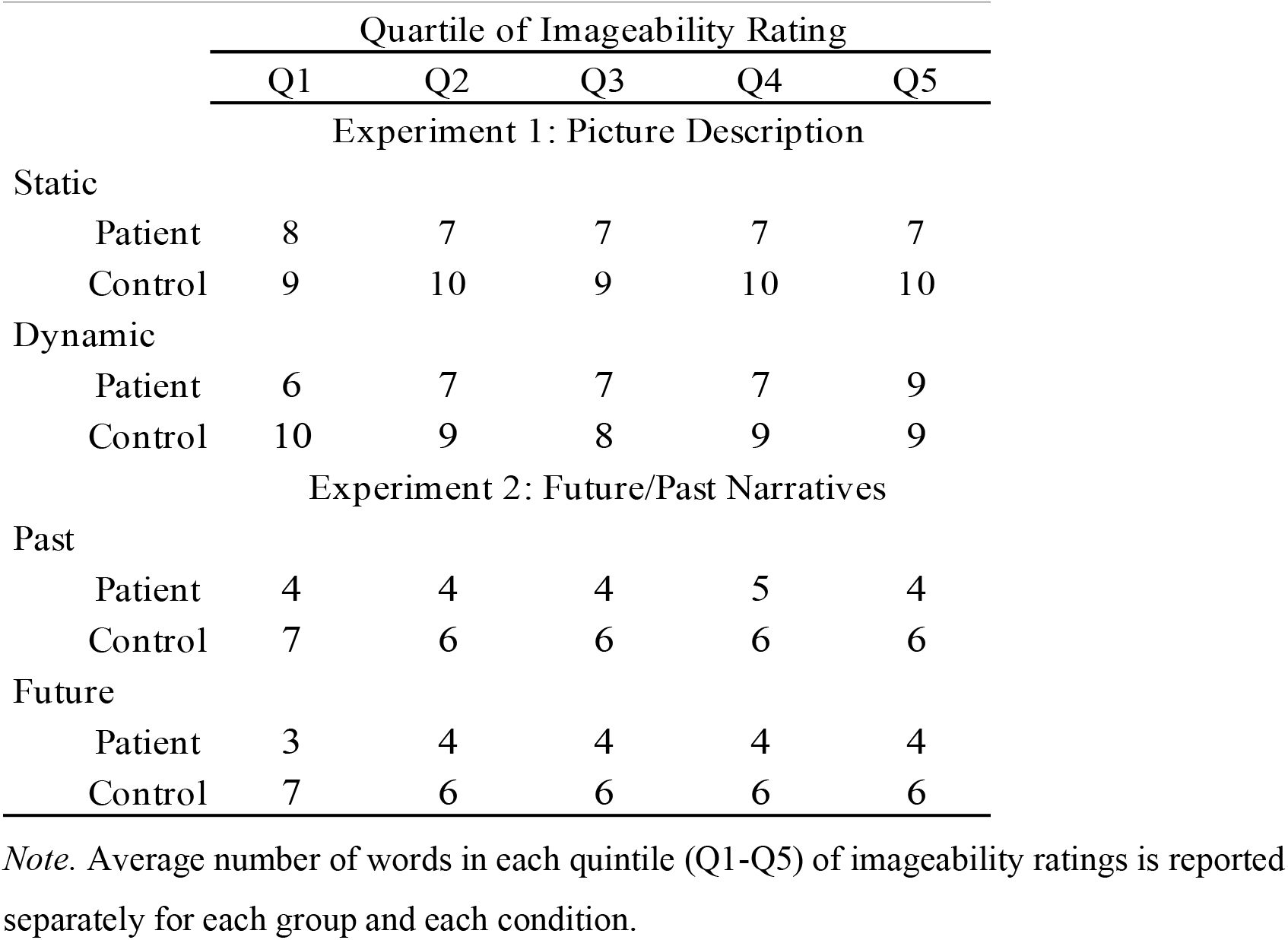
Distribution of Imageable Words in Experiment 1 and Experiment 2

